# ESM-LoRA-Gly: Improved prediction of N- and O-linked glycosylation sites by tuning protein language models with low-rank adaptation (LoRA)

**DOI:** 10.1101/2025.08.12.669850

**Authors:** Zhiyong Feng, Xing Zhang, He Wang, Xu Hong, Jian Zhan, Yaoqi Zhou

## Abstract

Glycosylation associates with many diseases ranging from cancer to neurodegeneration and understanding these disease mechanisms requires the precise identification of glycosylation sites. Computational prediction of glycosylation sites has been useful to complement laborious experimental methods, while existing tools lack sufficient accuracy and scalability. Here, we introduce ESM-LoRA-Gly, a method that employs Low-Rank Adaptation (LoRA) to fine-tune the ESM2-3B protein language model for predicting both N- and O-linked glycosylation sites. According to the evaluation on the benchmark datasets, ESM-LoRA-Gly outperforms existing state-of-the-art techniques. The improvement is particularly significant (>100% in Matthews correlation coefficient) for the O-linked dataset. By substantially reducing trainable parameters while maintaining predictive power, ESM-LoRA-Gly enables computationally efficient proteome-scale predictions. This approach should be instrumental for advancing glycoproteomic research and accelerating therapeutic discovery for glycosylation-related diseases.

## INTRODUCTION

Glycosylation, a crucial post-translational modification (PTM), involves the enzymatic attachment of sugar molecules, or glycans, to specific amino acid residues within proteins or lipids [1-2]. This process occurs widely in diverse species and cell types, and over half of human proteins are glycosylated [3-5]. It has two primary types, N-linked and O-linked glycosylation, which play distinct yet vital roles in cellular functions. N-linked glycosylation attaches oligosaccharides to asparagine residues, which occurs in the endoplasmic reticulum and is the basis for proper protein folding, stabilization, and trafficking [6-8]. O-linked glycosylation, however, involves the addition of N-acetylgalactosamine (N-GalNAc) to serine or threonine residues, which is critical for regulating protein activity and linking nutrient supply to protein function and stability. Dysregulation of either N-linked or O-linked glycosylation has been implicated in various diseases, including cancer, diabetes, and neurodegenerative disorders [9-10].

N-linked glycosylation sites are frequently identified using mass spectrometry [11-13], and considerable advancements have been made in experimental techniques for mapping and quantifying this post-translational modification. To date, over 14,000 unique N-linked glycosylation sites (N-glycosites) have been identified in humans [13]. Traditionally, O-linked glycosylation has been determined using chemical methods [14], such as β-elimination or hydrazine hydrolysis. Although experimental methods remain the most reliable ways to identify glycosites, these techniques can lead to peptide bone degradation and loss of glycan-specific information. Moreover, they are often time-consuming, labor-intensive, and limited in coverage. Consequently, a substantial portion of the proteome remains unannotated, underscoring the growing importance of complementary computational tools for the characterization of glycosylation sites.

Facilitated by machine learning methods, computational models have been developed to predict glycosylation sites. Net-NGlyc [15] was one of the early representatives that utilized an artificial neural network (ANN) for N-linked glycosylation prediction. EnsembleGly [16] employed an ensemble support vector machines (SVM) with input features of physicochemical properties, the position specific scoring matrix (PSSM) generated by PSI-BLAST [17], and the one-hot encoding. GlycoPP [18] also utilized an SVM, incorporating amino acid composition (AAC), one-hot encoding, PSSM, secondary structures (SS), and accessible surface area (ASA) features. NGlycPred [19] used a random forest classifier, leveraging sequence, pattern, and structure-based features to predict N-linked glycosylation sites. GlycoEP [20] and SPRINT-Gly [21] encoded data with sequence, evolutionary, and structural features for SVM and artificial neural networks, respectively, to identify N-linked glycosylation sites. OGP [22] was the model for predicting O-linked glycosylation, using a

Random Forest (RF) algorithm. More recently, deep learning techniques, particularly those leveraging protein language models, have emerged. For instance, DeepNGlyPred [23] employed a multilayer perceptron (MLP) restricted to N-X-[S/T] sequences, incorporating structural features including secondary structure, accessible surface area, relative solvent accessibility (RSA), torsion angles (Φ, Ψ), and disordered regions. LMNglyPred [24] utilized the protein language model ProtT5-XXL for sequence encoding, with an MLP to predict glycosylation sites. EMNGlyPred [25] applied the pre-trained protein language models (Evolutionary Scale Modeling, ESM) [26] and the pre-trained protein structure models (Inverse Folding Model) [27] for feature extraction, followed by SVM for glycosylation site prediction. HOTGpred [28] combined pre-trained protein language model embeddings and feature selection with Extreme Gradient Boosting (XGBT) to predict O-linked threonine glycosylation sites (OTGs).

Although language model-based predictors have significantly improved the accuracy of glycosylation prediction, they still face many limitations. For example, some predictors [23,24,28] simply used language model embeddings as one of the input features and the language model itself was not fine-tuned on the relevant sequences. As a result, the extracted embeddings cannot be perfectly adapted to the glycosylation prediction task. On the other hand, fine-tuning the language model with all parameters [29,30] on the glycosylation dataset not only requires substantial computational resources but also may risk fundamentally altering the model’s original capabilities and causing overfitting. Thus, a lightweight fine-tuning method is more desirable, one that can adjust language models to fit the glycosylation prediction task while avoiding breaking their inherent characteristics.

Recently, Low-Rank Adaptation (LoRA) [31] has emerged as an effective approach to address the challenges of fine-tuning large-scale pre-trained models. By decomposing model parameters into low-rank matrices, LoRA significantly reduces computational costs while maintaining model performance during fine-tuning. This efficient adaptation strategy has been successfully applied across various domains, including natural language processing, computer vision, and bioinformatics. Notably, Lv et al. [32] developed ProLLM, which utilizes LoRA to adapt general large language models (LLMs) for diverse protein language processing tasks. Similarly, Jeon et al. [33] proposed LoRA-BERT to enhance RNA sequence classification, demonstrating improved accuracy in distinguishing lncRNAs from mRNAs. In structural biology, Samuel Sledzieski et al. [34] employed LoRA for efficient fine-tuning of ESM2, achieving accurate protein-protein interaction predictions with minimal computational overhead. Further supporting its versatility, Robert Schmirler et al. [35] demonstrated that task-specific fine-tuning with LoRA enhances performance across various protein modeling tasks while maintaining computational efficiency. Building upon these advancements, we present ESM-LoRA-Gly, a novel method that leverages LoRA to fine-tune the ESM-3B protein language model. Results confirmed ESM-LoRA-Gly as a model with a reduced training parameters while having superior performance in prediction of N- and O-linked glycosylation sites, when compared to existing tools.

## MATERIAL AND METHODS

### Dataset for N-linked glycosylation prediction

For N-linked glycosylation prediction, we integrated multiple datasets to ensure comprehensive coverage and enhance model generalization. The primary dataset comprised 12,534 positively annotated sites from DeepNGlyPred (sourced from NGlycositeAtlas [36]), along with an equal number of negative samples (non-glycosylation sites) from DeepLoc-1.0. To further expand the training data, we incorporated benchmark datasets from SPRINT-gly, PUstackgly, and EMNGly to build a combined collection of 11,083 glycoproteins. After redundancy removal and sequence filtering (restricted to the N-X-[S/T] motif), the final dataset retained 14,869 positive glycosylation sites and 12,533 negative samples.

To avoid overfitting, we employed CD-HIT [37] to eliminate proteins with sequence identities exceeding 30%, yielding a refined dataset of 12,326 positive sites and 11,848 negative sites across 9,059 proteins. The dataset was first divided into training and test sets at a 9:1 ratio using a protein sequence-based splitting strategy, with no identical proteins in the two datasets. Subsequently, the training set was further split into training and validation subsets using the same 9:1 ratio, while maintaining strict protein separation. It should be emphasized that all glycosylation sites from the same protein sequence were exclusively assigned to a single partition (training, validation, or test set), preventing overlap across these sets.

### Dataset for O-linked glycosylation prediction

We obtained the O-linked glycosylation site dataset from the O-Glycoprotein (OGP) repository [2222], a comprehensive database of experimentally characterized O-glycoproteins developed by Huang et al. in 2021. The dataset contained 2,403 protein sequences with 11,813 positive sites. To minimize redundancy, we applied CD-HIT to remove sequences with >30% similarity, resulting in 1,397 proteins and 11,546 positive sites.

Following the same partitioning strategy used for the N-linked datasets, we randomly split this non-redundant dataset into training and test sets at a 9:1 ratio. The training set included 1,256 proteins, 10,537 positive sites, and 123,774 negative sites, while the test set comprised 141 proteins, 1,009 positive sites, and 14,645 negative sites. To address class imbalance, we performed downsampling to equalize positive and negative samples. The training set was further divided into training and validation subsets at a 9:1 ratio. After balancing, the training set contained 1,130 proteins, 9,946 positive sites, and 9,470 negative sites; the validation set had 126 proteins, 591 positive sites, and 591 negative sites; and the independent test set included 141 proteins, 1,009 positive sites, and 1,009 negative sites. As for N-linked datasets, we ensure that all glycosylation sites in a single protein are only allowed in a single dataset (training, validation, or test set). Detailed statistics are provided in Table 1.

**Table 1.**
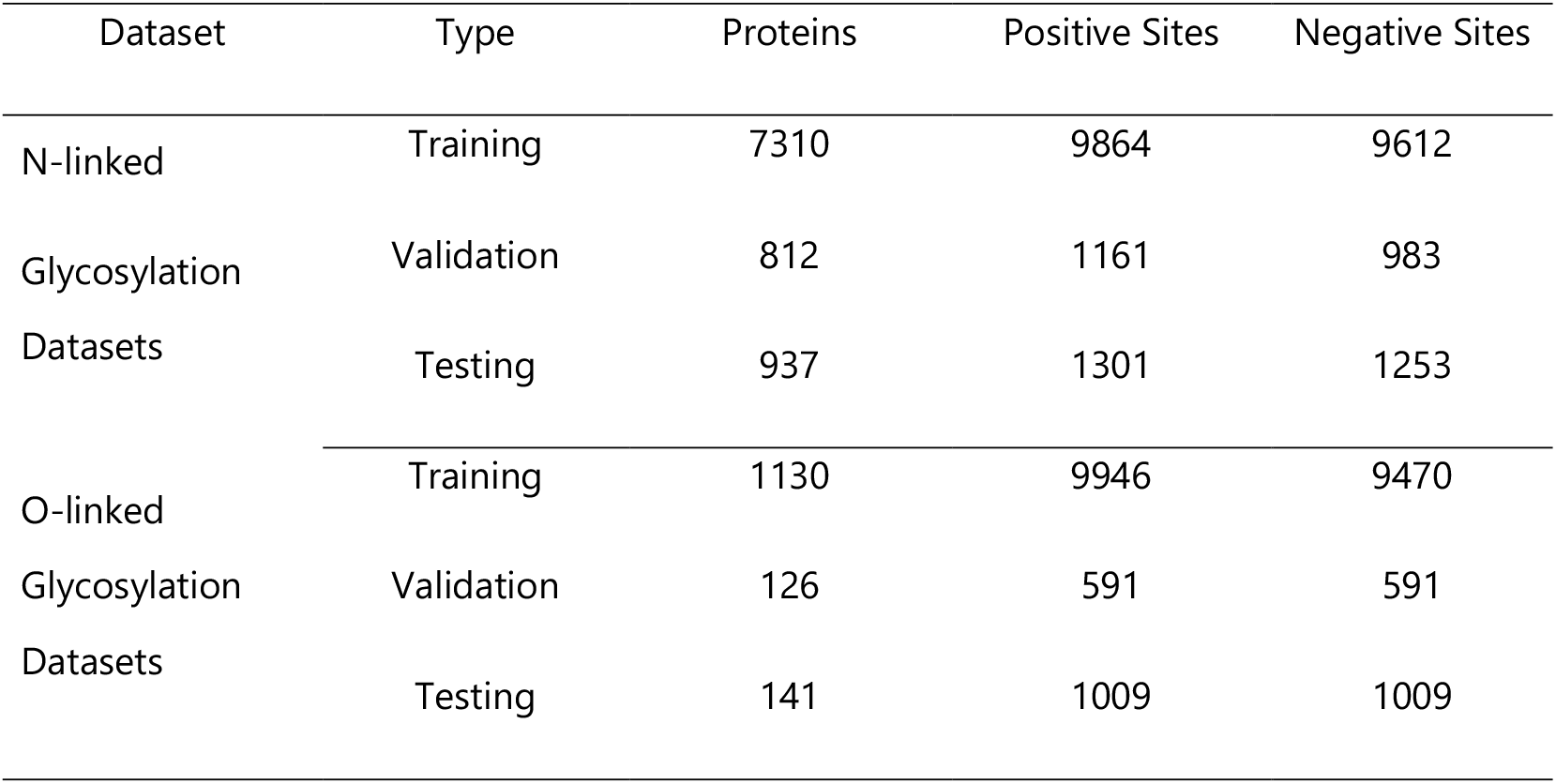
Statistics of positive and negative sites for training, validation, and testing for N-linked and O-linked glycosylations.

### Network Architecture

The overall architecture of our model is depicted in Figure 1. Specifically, our method leverages ESM-2 [38] and LoRA for efficient fine-tuning of glycosylation site prediction. As shown in Figure 1A, input protein sequences are first embedded using ESM-2. These embeddings are processed through the 36-layer ESM-2 encoder. During fine-tuning, LoRA modules are incorporated into the query (*W*_*q’*_), key (*W*_*k*_), value (*W*_*v*_), and output linear layers (*W*_*o*_) within each attention block of the encoder, as detailed in Figure 1B. The encoder output is then fed through a series of hidden layers to a single Multilayer Perceptron (MLP). This MLP serves as the classification head, performing binary classification to distinguish glycosylation sites from non-glycosylation sites. For supervised training, LoRA refines only the attention layer parameters (*W*_*q*,_ *W*_*k‘*_ *W*_*v*_) and output layer of the encoder (*W*_*o*_) by decomposing weight updates Δ*W* into low-rank matrices *A* and *B*, whereΔ*W* ∈ *R*^*d×*r^, *A* ∈ *R*^*d×h*^, and *B* ∈ *R*^*h×*r^ with rank r=8. Based on the evaluation of ESM2 models of various sizes, ESM-LoRA-Gly employed the ESM2-3B architecture (3 billion parameters) as the final model.

**Figure 1:**
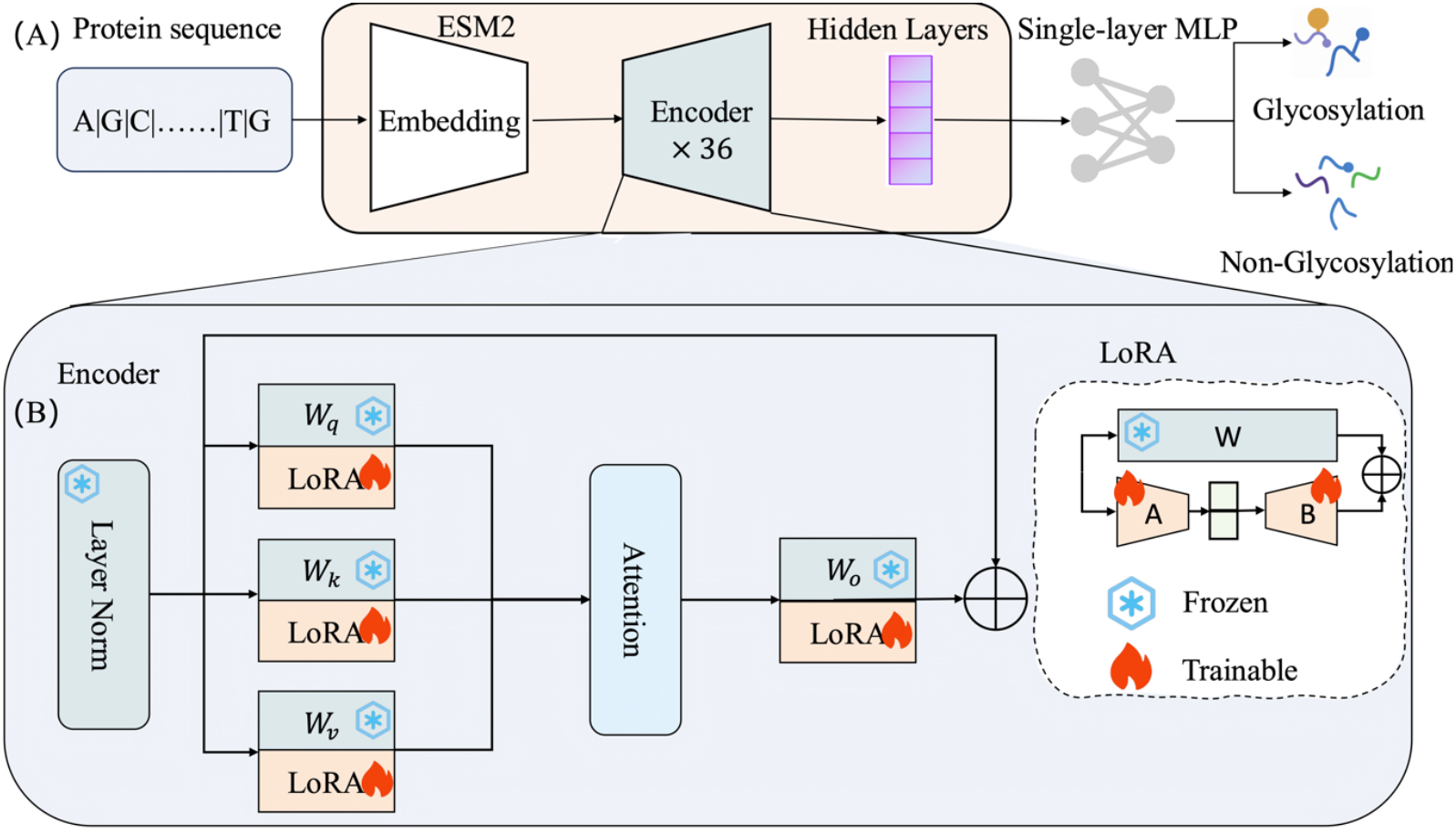
(A) The overall architecture of our model: sequence embeddings are generated from ESM-2 with an input of query protein sequence, followed by utilizing the Low-rank adapters (LoRA) encoder to train an MLP for prediction of glycosylation sites. (B) The detailed architecture for training the LoRA encoder, *W*_*q’*_ *W*_*k‘*_ *W*_*v*_ are the parameters of the ESM2 encoder attention layer query, key, and value, and *W*_*o*_ is the output layer of the encoder. LoRA refines *W*_*q’*_ *W*_*k‘*_ *W*_*v*_ by decomposing the changes *ΔW* as the multiplication product of two matrices *A* and *B* of lower rank *r* (*ΔW* = *AB*) and minimizing *A* and *B* with fewer parameters.

### Evaluation Metrics

To evaluate model performance, we employed four metrics: accuracy (ACC), sensitivity (SN), specificity (SP), and Matthews Correlation Coefficient (MCC), defined as follows:

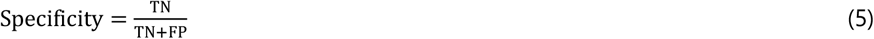

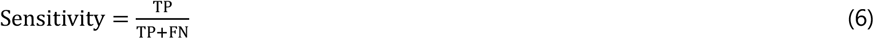

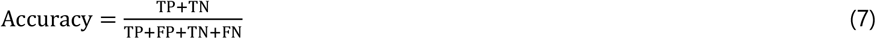

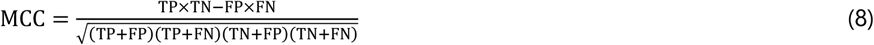

where Specificity (SP), sensitivity (SN), and accuracy (ACC) range from 0 to 1, typically expressed as percentages. The Matthews Correlation Coefficient (MCC) ranges from -1 to 1, where 1 indicates perfect prediction, 0 indicates random guessing, and -1 indicates complete inverse prediction relative to true labels. MCC is particularly valuable for imbalanced datasets as it incorporates all four confusion matrix elements: true positives (TP), true negatives (TN), false positives (FP), and false negatives (FN).

### Experimental Setup

Our experimental setup comprised both software and hardware components. The software stack utilized Python 3.9, the transformers library (v4.33), and the peft library (v0.11.0). The hardware configuration included four NVIDIA A800-SXM4-80GB GPUs. Hyperparameters and LoRA configurations are specified in Table 2 and Table 3, respectively.

**Table 2.**
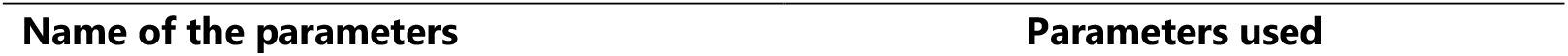
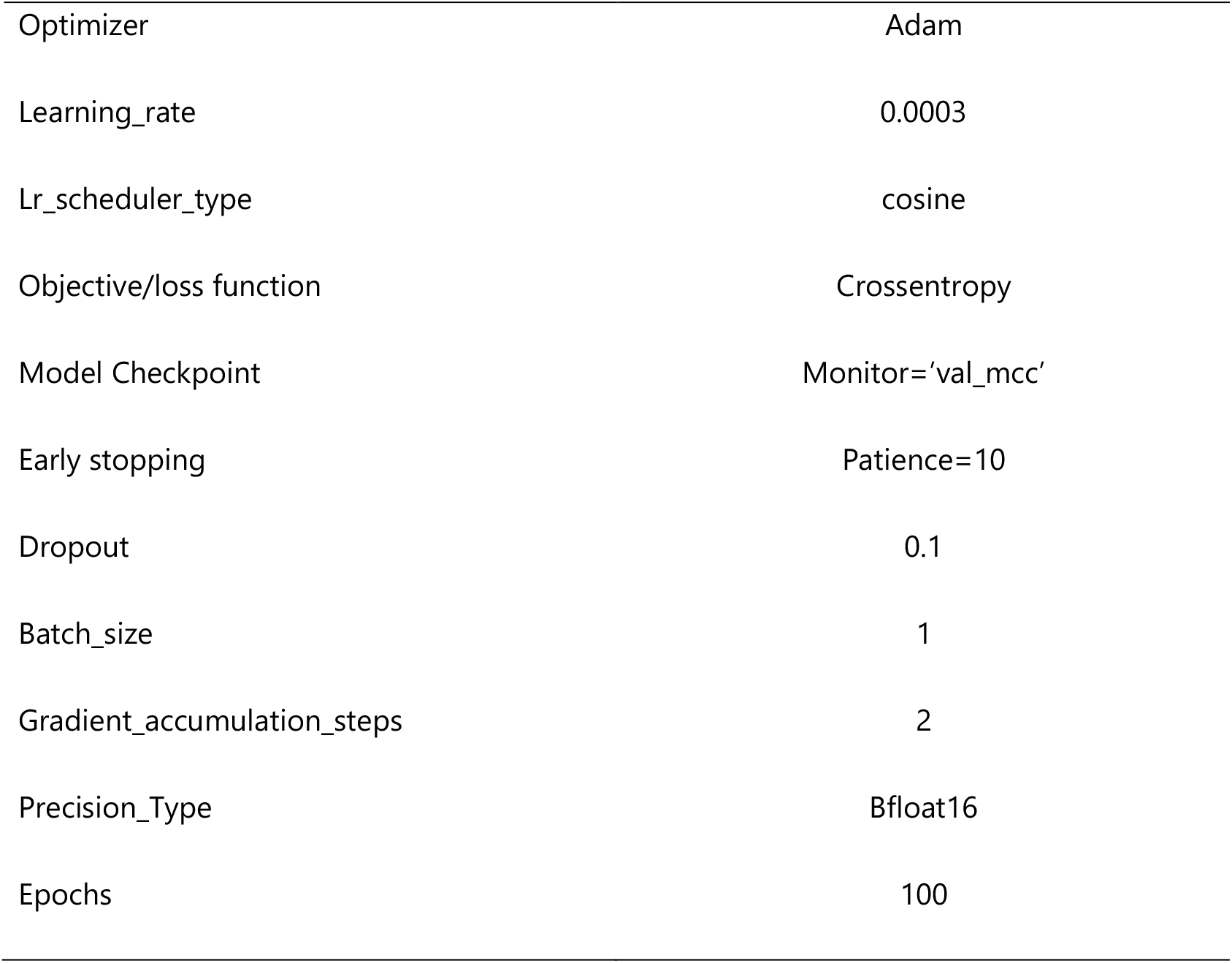
Hyperparameters employed in the model.

| Name of the parameters | Parameters used |
| --- | --- |
| Optimizer | Adam |
| Learning_rate | 0.0003 |
| Lr_scheduler_type | cosine |
| Objective/loss function | Crossentropy |
| Model Checkpoint | Monitor='val_mcc' |
| Early stopping | Patience=10 |
| Dropout | 0.1 |
| Batch_size | 1 |
| Gradient_accumulation_steps | 2 |
| Precision_Type | Bfloat16 |
| Epochs | 100 |

**Table 3.** The LoRA config for the model.

| Name of the parameters | Parameters used |
| --- | --- |
| Task_type | SEQ_CLS |
| inference_mode | False |
| r | 8 |
| lora_alpha | 32 |
| lora_dropout | 0.1 |
| bias | 'all' |
| target_modules | ['query', 'key', 'value', 'out proj'] |

### Methods Compared

The results of HOTGpred [28] and OGP [22] were obtained from the webservers: https://balalab-skku.org/HOTGpred/ and http://www.oglyp.org/predict.php, respectively.

LMNGlyPred [24] and EMNGly [25]: We obtain the code and weights from https://github.com/KCLabMTU/LMNglyPred and https://github.com/StellaHxy/EMNgly/tree/master, respectively. They were recompiled according to the author’s operating environment.

## RESULTS

### Performance on the N-linked Dataset

To evaluate the predictive performance of our model, ESM-LoRA-Gly was compared with two recent language model-based predictors, LMNGlyPred and EMNGly, on the N-linked glycosylation dataset. As summarized in Table 4 and illustrated in Figure 2, ESM-LoRA-Gly exhibits superior performance across all standard evaluation metrics. The notably high Area Under the Receiver Operating Characteristic Curve (AUC) and Area Under the Precision-Recall Curve (AUC-PR) values are particularly significant, highlighting the model’s exceptional discriminative capability and effectiveness on imbalanced datasets, respectively.

**Table 4.** Performance comparison of ESM-LoRA-Gly with other N-linked glycosylation site predictors on the test set, evaluated using MCC, accuracy, specificity, sensitivity, AUC-ROC, and AUC-PR.

| Model | MCC | Accuracy | Specificity | Sensitivity | AUC | AUC-PR |
| --- | --- | --- | --- | --- | --- | --- |
| LMNGlyPred | 0.494 | 0.757 | 0.753 | 0.763 | 0.828 | 0.687 |
| EMNGly | 0.828 | 0.914 | 0.924 | 0.904 | 0.976 | 0.976 |
| ESM-LoRA-Gly | **0.868** | **0.934** | **0.953** | **0.915** | **0.981** | **0.983** |

**Figure 2.**
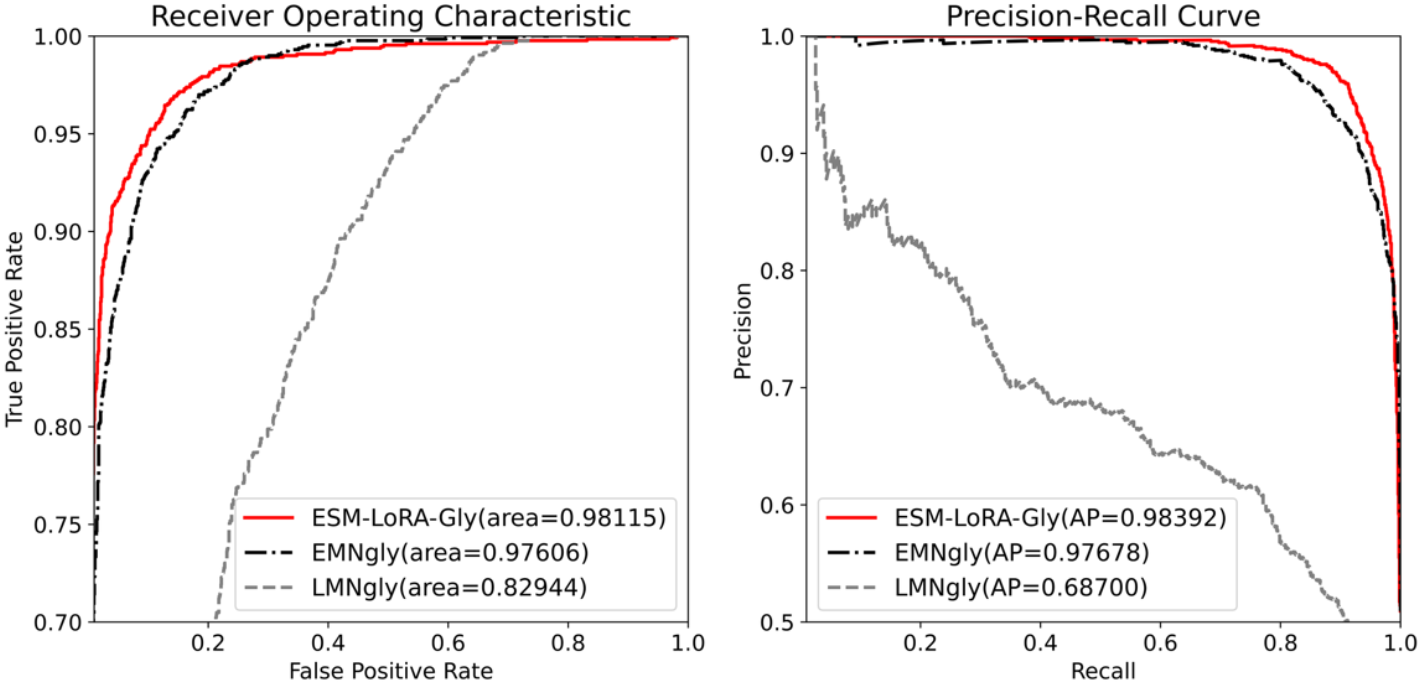
Performance comparison in the independent test set of the N-linked dataset among EMNgly, LMNgly, and the proposed method ESM-LoRA-Gly.

Quantitatively, ESM-LoRA-Gly achieves an AUC-PR of 0.983 and a Matthews Correlation Coefficient (MCC) of 0.868. This MCC represents a substantial improvement of ∼5% over the model EMNGly (0.828) and a 76% enhancement over LMNGlyPred (0.494). These results demonstrate the effectiveness of our proposed fine-tuning approach.

### Performance on the O-linked Dataset

For O-linked glycosylation prediction, we compared ESM-LoRA-Gly with diverse existing predictors: a traditional machine learning method (Random Forest), OGP, and a recent protein language model-based approach, HOTGpred. As detailed in Figure 3 and Table 5, ESM-LoRA-Gly achieves superior performance across all evaluation metrics.

**Table 5.**
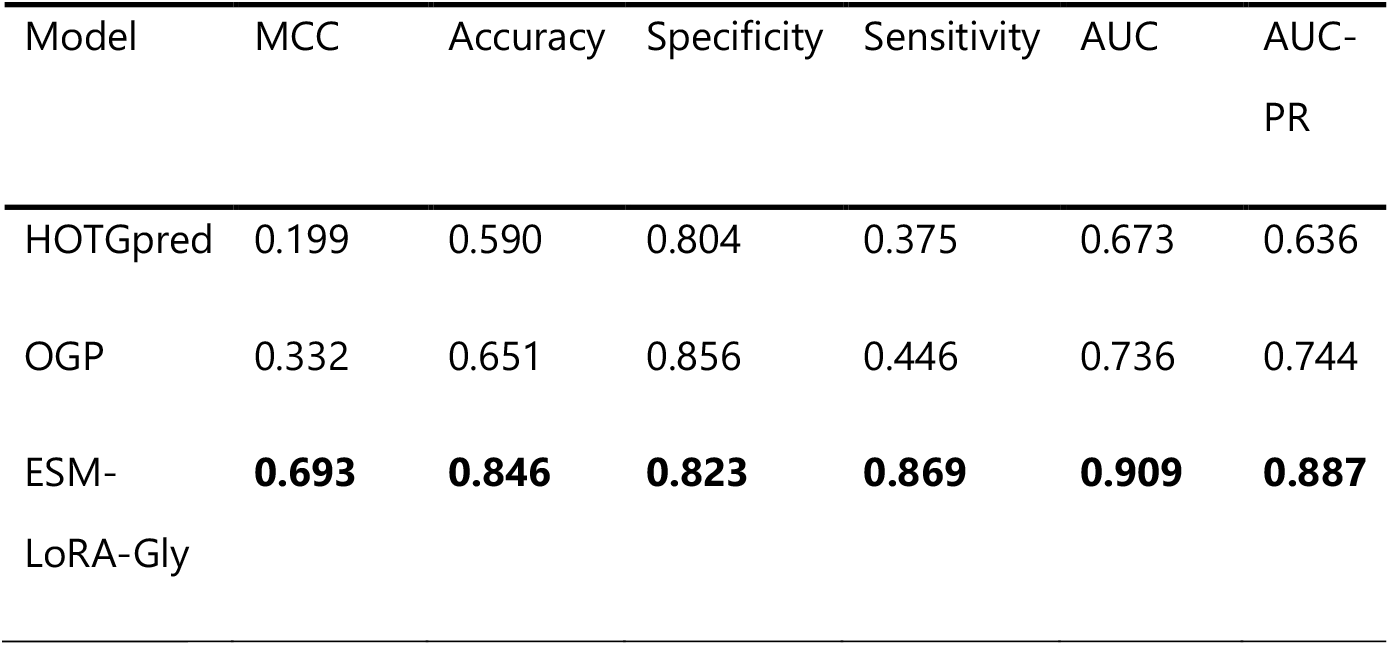
Performance comparison of ESM-LoRA-Gly with other glycosylation site predictors on the O-linked dataset, evaluated using MCC, accuracy, specificity, sensitivity, AUC-ROC, and AUC-PR.

| Model | MCC | Accuracy | Specificity | Sensitivity | AUC | AUC-PR |
| --- | --- | --- | --- | --- | --- | --- |
| HOTGpred | 0.199 | 0.590 | 0.804 | 0.375 | 0.673 | 0.636 |
| OGP | 0.332 | 0.651 | 0.856 | 0.446 | 0.736 | 0.744 |
| ESM-LoRA-Gly | **0.693** | **0.846** | **0.823** | **0.869** | **0.909** | **0.887** |

**Figure 3.**
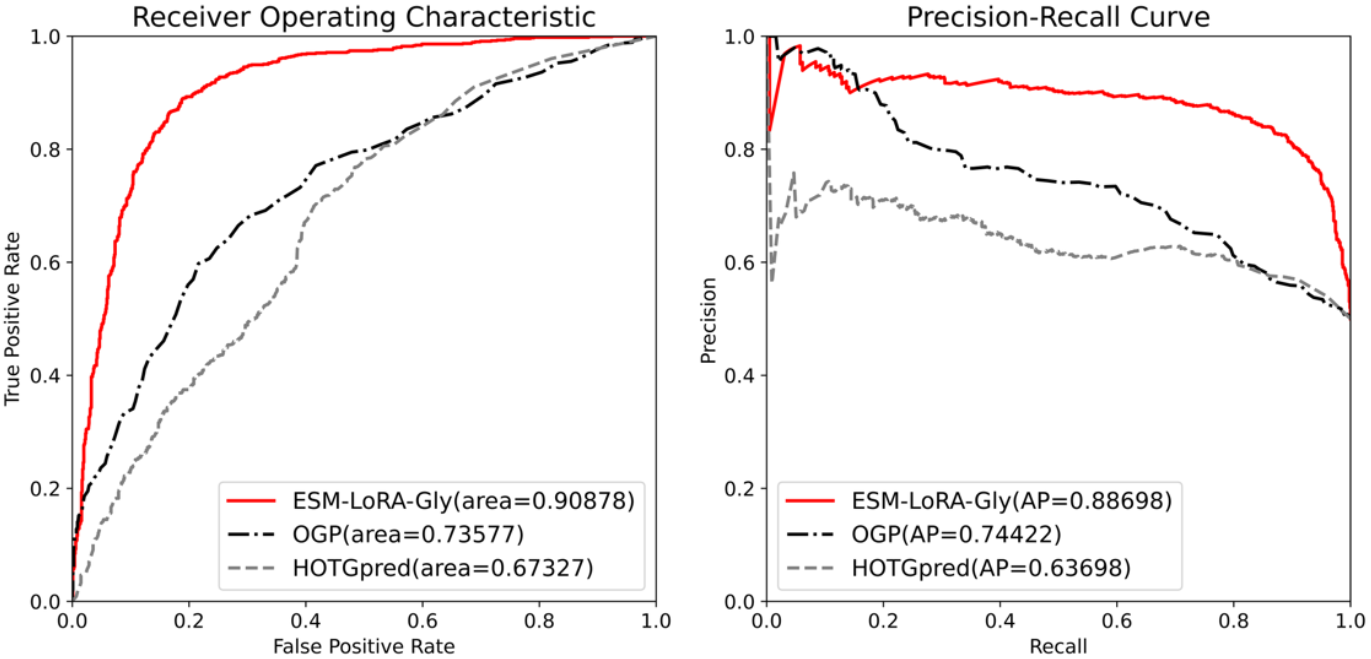
Performance comparison on the independent test set of the O-linked dataset among the proposed ESM-LoRA-Gly model, OGP, and HOTGpred.

Beyond its enhanced accuracy, ESM-LoRA-Gly attains an AUC-PR score of 0.887, demonstrating robust predictive capability. The most notable improvement is observed in the Matthews Correlation Coefficient (MCC), where ESM-LoRA-Gly achieves 0.693, significantly exceeding the second-best method, OGP (0.332), by 108%. This substantial enhancement highlights the ability of our fine-tuned ESM-2 model to identify complex sequence patterns that govern O-linked glycosylation.

### Robustness of the performance according to 10-fold cross-validation

To assess performance stability, we performed 10-fold cross-validation independently on the N-linked and O-linked datasets. Each dataset was randomly divided into 10 equal subsets. In each iteration, one subset served as the validation set, and the remaining nine subsets were employed for model training. This process was repeated 10 times, ensuring that every subset was validated exactly once. The prediction metrics (e.g., MCC) from all iterations were aggregated to compute the mean and standard deviation presented in Table 6. As the Table shows, the low standard deviation values for all metrics measuring the performance on N- and O-linked datasets demonstrate the method’s robustness against variations in data partitioning.

**Table 6.**
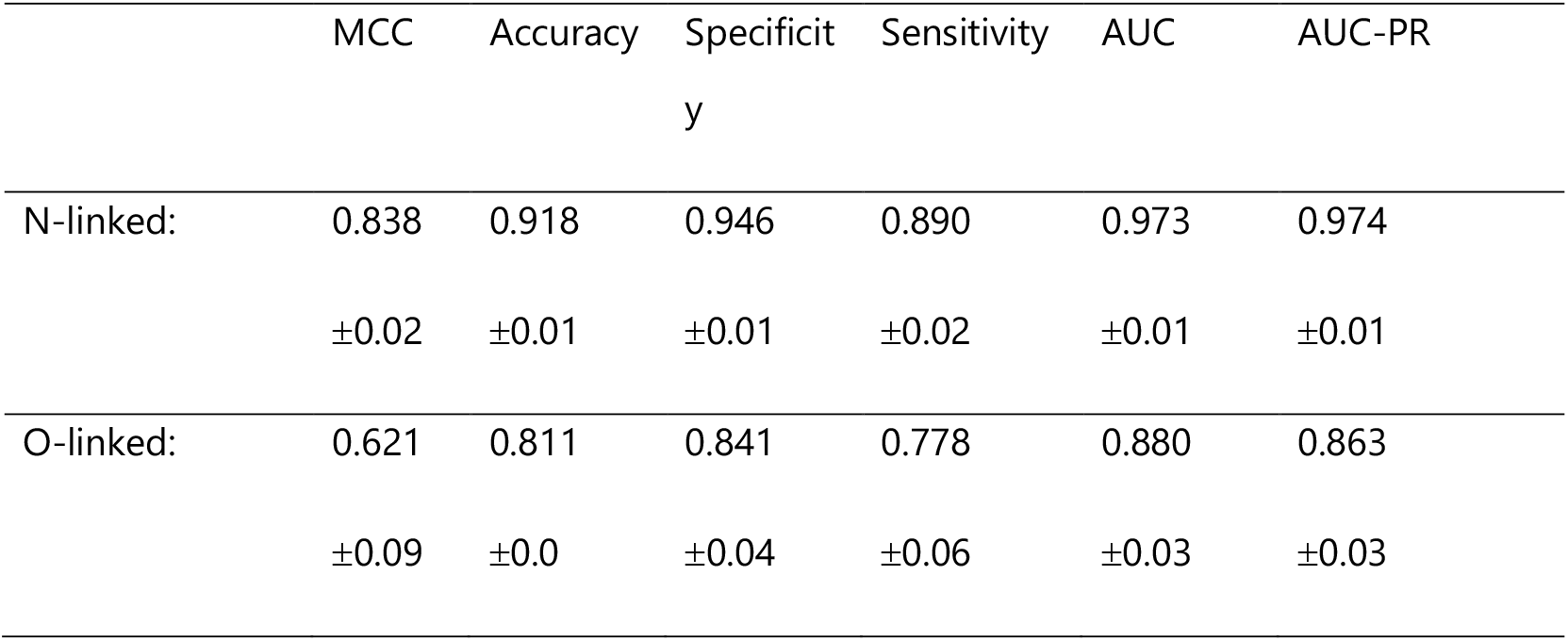
Mean and standard deviation of prediction metrics from 10-fold cross-validation on the N-linked and O-linked datasets.

### Ablation study

We conducted ablation studies on the N-linked dataset to evaluate LoRA’s effectiveness under different parameter configurations and assess the impact of pre-trained model scale on performance. As presented in Table 7 and Figure 4, three training paradigms were compared: linear regression (training only the fully connected layer), full-parameter fine-tuning, and LoRA. For linear regression, the ESM2-3B-based model achieved the highest MCC, AUC, and accuracy. For full-parameter fine-tuning—limited to ESM2-650M due to computational constraints—this approach outperformed linear regression using the same ESM2-650M backbone. Conversely, LoRA attained optimal performance with ESM2-3B. Notably, when applied to ESM2-650M, LoRA (MCC=0.844) surpassed both full fine-tuning (MCC=0.835) and linear regression (MCC=0.812), demonstrating that parameter expansion is not essential for the best results. Collectively, these findings reveal that state-of-the-art performance arises from synergistically combining the representational capacity of large-scale models (ESM2-3B) with parameter-efficient adaptation (LoRA).

**Table 7.** Prediction performance of different models and methods on the N-linked dataset.

| Method | Model | MCC | Accuracy | Specificity | Sensitivity | AUC |
| --- | --- | --- | --- | --- | --- | --- |
| Linear<br>Regression | ESM2-8M | 0.724 | 0.860 | 0.887 | 0.837 | 0.939 |
|  | ESM2-35M | 0.781 | 0.890 | 0.861 | 0.918 | 0.963 |
|  | ESM2-150M | 0.842 | 0.919 | 0.961 | 0.879 | 0.968 |
|  | ESM2-650M | 0.812 | 0.900 | 0.93 | 0.87 | 0.971 |
|  | ESM2-3B | 0.830 | 0.915 | 0.908 | 0.923 | 0.973 |
|  | ESM2-15B | 0.820 | 0.909 | 0.941 | 0.877 | 0.972 |
| Finetune | ESM2-8M | 0.800 | 0.900 | 0.892 | 0.909 | 0.967 |
|  | ESM2-35M | 0.791 | 0.895 | 0.904 | 0.887 | 0.958 |
|  | ESM2-150M | 0.837 | 0.917 | 0.951 | 0.885 | 0.950 |
|  | ESM2-650M | 0.835 | 0.918 | 0.928 | 0.901 | 0.974 |
| LoRA | ESM2-8M | 0.815 | 0.900 | 0.925 | 0.868 | 0.971 |
|  | ESM2-35M | 0.825 | 0.911 | 0.954 | 0.869 | 0.968 |
|  | ESM2-150M | 0.848 | 0.923 | 0.946 | 0.901 | 0.977 |
|  | ESM2-650M | 0.844 | 0.916 | 0.937 | 0.899 | 0.976 |
|  | **ESM2-3B** | **0.868** | **0.934** | **0.953** | **0.915** | **0.981** |
|  | ESM2-15B | 0.832 | 0.916 | 0.911 | 0.922 | 0.972 |

**Figure 4.**
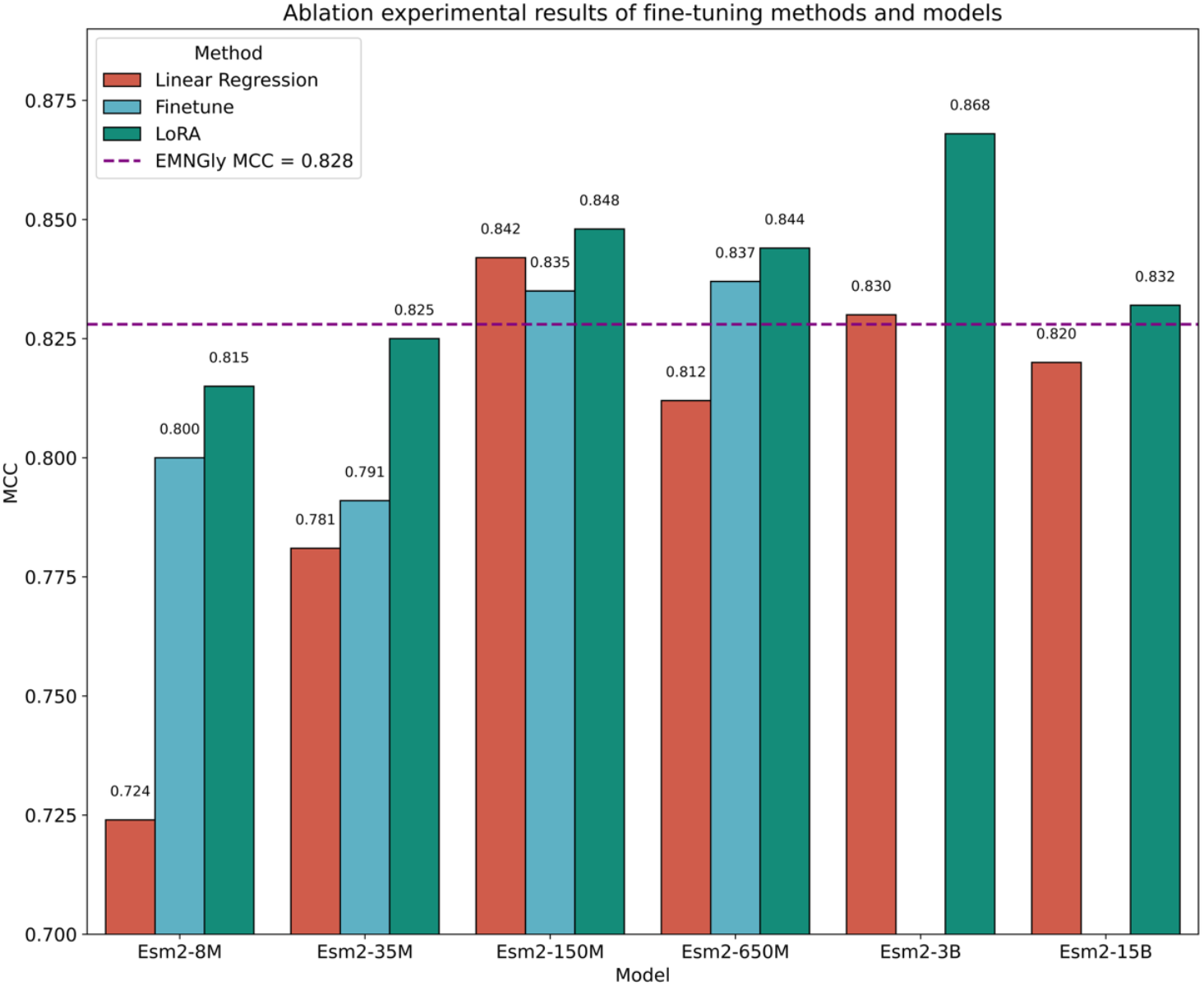
Prediction performance of ablation studies on the N-linked dataset.

## DISCUSSION

The application of LoRA for fine-tuning large pre-trained protein language models constitutes a significant advancement in task-specific model refinement. Here, we utilize LoRA for glycosylation site prediction, achieving superior performance while maintaining computational efficiency in both training and inference, thereby providing a scalable solution for large-scale proteomics studies. For instance, during ESM-3B training, LoRA requires an average memory usage of 200GB, substantially lower than the 280GB consumed by full fine-tuning or linear regression. Inference time is approximately 1 minute per protein (1000 amino acids) on a system equipped with single NVIDIA A100-80GB GPUs. Consequently, ESM-LoRA-Gly facilitates feasible proteomics-scale predictions.

In addition to LoRA, there are other fine-tuning techniques such as AdaLoRA [3939] and IA3 [40]. AdaLoRA [39] extends LoRA by dynamically allocating low-rank matrices during training and automatically adjusting layer-wise rank allocation to enhance parameter efficiency.

However, this dynamic adjustment causes continuous weight matrix shape changes, impeding tensor synchronization across multiple GPUs and limiting practical efficiency. Computational constraints precluded AdaLoRA training on our dataset. IA3 [40] fundamentally differs from LoRA in adaptation: while LoRA adds fixed low-rank matrices to pretrained weights, IA3 applies learnable scaling vectors directly to intermediate activations within attention and feed-forward layers. This approach yields fewer trainable parameters and lower computational overhead but may restrict expressiveness compared to LoRA’s direct weight modulation. When applied to ESM2-3B for glycosylation site prediction, IA3 exhibited markedly poorer performance (MCC =0.780) relative to LoRA (MCC=0.868).

Data and Source Code Availability. All data and software codes, along with trained parameters, are available at https://github.com/NancyFyong/ESM-LoRA-Gly.

## ACKNOWLEDGEMENTS

We gratefully acknowledge the High Performance Computing Cluster at Shenzhen Bay Laboratory and Shenzhen Medical Academy of Research and Translation for completing this study.

